# Pyrethroid and metal residues in different coffee bean preparing processes and their human health risk assessments via consumption

**DOI:** 10.1101/2021.03.25.436952

**Authors:** Paphatsara Khunlert, Phanwimol Tanhan, Amnart Poapolathep, Saranya Poapolathep, Kanjana Imsilp

## Abstract

The study was conducted on 50 samples of coffee beans from various origins. The samples included green coffee beans, roasted beans, brew coffee drinks and coffee sludge. Three processes were used to prepare these samples: dried, semi-washed, and washed. Three synthetic pyrethroid insecticides and nine heavy metals were subsequently analyzed using modified Quick, Easy, Cheap, Effective, Rugged and Safe (QuEChERS) and acid digestion methods, respectively. The quantification of pyrethroids was performed by GC-μECD whereas those of metals were determined using flame atomic absorption spectrophotometer. According to the results, concentrations of both pyrethroids and heavy metals were predominantly found in green coffee beans except for Cr. Pyrethroid insecticides were not detectable in brew coffee drink and heavy metal concentrations were below the acceptable daily intake (ADI) level. Risk estimations for daily coffee intake using the health risk indices (HRIs) and target hazard quotients (THQs) of normal and the 97.5 percentile Thai consumers were less than 1. This indicated that the coffee drinks from studied samples could not cause potential health risk.

## Introduction

Coffee is a worldwide favorite beverage. It has several beneficial antioxidants and is one among the richest chlorogenic acid sources (1). However, coffee drinking can potentially lead human to expose to multiple toxicants. Those of which include pesticides, heavy metals, organic solvents and pharmaceutical agents. These hazardous chemicals have abundantly been used to protect coffee trees from pests and diseases during cultivation and manufacturing processes (2). Even though coffee beans are roasted, pesticide residues are still present. Recent studies revealed that coffee beans and coffee wastes were contaminated by numerous pesticides and metals (3–5).

Applications of pesticides and insecticides to coffee crops are either directly to soil or to aerial parts of the plants (6). Chemical classes generally used are organophosphates, pyrethorids, carbamates, chlorinated cyclodienes and organotins (6). The most favorite class among those are pyrethroids and accounted for one fifth of total insecticide global market (7, 8). They are synthetic analogues of pyrethrin found in *Tanacetum cinerariaefolium* (Asteraceae) or pyrethrum daisy. Pyrethroids have extensively been used in agricultural crops as well as in animals to reduce pests and decrease their transfers which can help improving the coffee yield. Pyrethroids commonly used in coffee culture are cypermethrin, esfenvaleratem fenpropathrin, permethrin and cyfluthrin (9). Human can be exposed to these pyrethroids via inhalation, dermal contact, and ingestion (the major route) (10). The exposure can deteriorate human health. Even though harmful health effects following coffee consumption are likely, data are still limited.

Another group of toxicants that can be found in coffee beans is metals. They are naturally occurred or resulted from anthropogenic activities. The metals, either trace or toxic, have widely been dispersed in the environment and entered food chains at different concentrations. Coffee plants can absorb these metals through their roots and translocate them into the shoots as well as grains. Grains or beans usually accumulate metals at lower concentration than other parts. However, lifetime consumption of contaminated beans may pose adverse effects on human health. Coffee drinks commonly contain essential elements such as copper (Cu), manganese (Mn) and iron (Fe) at low concentrations. However, the essential [e.g., Cu, chromium (Cr), Mn, nickel (Ni), zinc (Zn)] and toxic [e.g., arsenic (As), cadmium (Cd), lead (Pb), cobalt (Co)] elements can be harmful if their levels exceed human tolerable limits.

Commonly, coffee is traded as green coffee beans prepared by either dry or washed processes. Both processes have similar steps from cleaning, separation, drying, storage, processing to classification. The differences are that, in the dry process, coffee fruits are sun dried followed by mechanical removal of dried husk whereas in the washed process, the damage and unripe coffee fruits are separated from the ripe ones by floatation technique. The skin and pulp in washed process were mechanically removed by pressing coffee fruits into water through metal sieve. Another method containing fewer steps is a semi-washed process. In this process, the outer skins are mechanically removed, and coffee beans are washed then dried. The quality of coffee beans in washed process generally scores higher than dried process coffee. This is because washed coffee beans have a higher percentage of ripe fruit harvested, while dry coffee beans have a wider range of ripeness from unripe to overripe. These also resulted in measurably different effects of sugar and flavor precursors present (11).

Several studies reported the contents of elements including heavy metals in green (raw) and roasted coffee varieties in different parts of the world (12) using a variety of techniques. The elements present in the roasted and ground coffee samples and their infusions can differ (9, 13). In addition, brewing types and roasting conditions also affected the concentration of elements in resulting infusions (14, 15). de Queiroz, Azevedo (9) determined heavy metals (Cd, Cr, Cu, Mn, Ni, Pb, Zn) in the roasted and ground coffee beans and brew. They reported that for all the infusions, the metals evaluated were found in lower concentrations with respect to the maximum permissible daily intake, except for Pb. Hence, it is important to determine heavy metal concentrations in various forms of coffee (raw, roasted, brew).

To the best of our knowledge, no research study has been conducted for analyzing the heavy metals and pyrethroid insecticides in green coffee beans, roasted coffee, and brew in Thailand. Therefore, the main goals of this study were 1) to determine the residues of pyrethroid insecticides (flumethrin, cypermethrin, and cyfluthrin) and heavy metals (Cd, Co, Cr, Cu, Fe, Mn, Ni, Pb, and Zn) in various forms of coffee (green, roasted, brew, sludge) originated from green coffee beans prepared by 3 processes (dry, semi-washed, washed); 2) to evaluate the human health risk from insecticide and metal contaminated coffee drink using the estimate daily intake/EDI, health risk index/HRI, and target hazard quotients/THQs. The results of this study would provide useful baseline data on the levels of contaminants and human health risks through coffee consumption.

## Materials and Methods

### Chemicals and reagents

All chemicals and reagents used in this study were ≥ 99% purity analytical or pesticide grades. Acetonitrile, ethyl acetate, acetic acid and nitric acid were purchased from RCI Labscan, Ltd., Thailand. Magnesium sulfate anhydrous (MgSO_4_; 98.0%), sodium chloride (NaCl), sodium acetate anhydrous (C_2_H_3_NaO_2_), tri-sodium citrate dihydrate (C_6_H_9_Na_3_O_9_), sodium citrate anhydrate (C_6_H_5_Na_3_O_7_) and alumina were purchased from Sigma-Aldrich, USA. Solid phase extractants including primary secondary amine (PSA) and octadecyl carbon (C_18_) were purchased from Agela Technologies, China and Macherey-Nagel, USA, respectively. Pyrethroid insecticide standards including cyfluthrin (4 isomers), flumethrin and cypermethrin (4 isomers) and individual metal standard solutions (Cd, Co, Cr, Cu, Fe, Mn, Ni, Pb and Zn; 1000 mg/L each) were purchased from Supelco^®^ Sigma-Aldrich, USA.

### Sample collection and preparation

Fifty 1-kg green coffee bean samples were randomly purchased from local markets in Bangkok, Thailand. They were from Asia, North America, South America, and South Africa. All samples were divided by their preparation processes into three groups; i.e., dried, semi-washed, and washed (Table 1). Each sample was aliquoted into two equal sub-samples. One sub-sample was ground and homogenized with miller and served as green coffee bean samples. Another sub-sample was roasted at 230°C until it presented the aroma (3). The roasted beans were ground into fine powder with a blender and 10 g were collected and added with 180 mL hot water (96°C). The brew coffee drink was then collected using paper drip technique. Coffee sludge remained in the filtered paper was also collected and oven dried at 65°C before storage. Green coffee beans from organic cultivation in Thailand were used as a blank and for spiked samples. They were prepared using similar processes as those of the samples. All samples were kept at −20°C until chemical analysis.

**Table 1.** Description of three coffee bean processes.

| Process | Description |
| --- | --- |
| Dried (D) | The rip beans were removed after sundrying coffee fruits. |
| Semi-washed (SW) | The hybrid method combining washed and dried processes. |
| Washed (W) | The rip beans were removed after coffee fruit fermentation. |

### Sample extraction and clean-up

#### Pyrethroid residues

To assess the differences in pyrethroid insecticide residues among coffee bean processes, the extraction and clean-up of green coffee beans and processed coffee samples (roasted coffee powder, brew coffee drink and coffee sludge) were conducted using modified Quick, Easy, Cheap, Effective, Rugged and Safe (QuEChERS) method combined with dispersive solid phase extraction (d-SPE) clean-up method. The extraction and clean-up procedures for green coffee beans and processed coffee samples were as follows:

##### Green coffee beans and brew coffee drink

2.5 g green coffee beans or 10 mL brew coffee drink were added with 10 mL acetonitrile, and hand shaken for 1 min. The samples were then added with 4 g MgSO_4_ and 1.5 g C_2_H_3_NaO_2_, and shaken for 1 min. They were subsequently centrifuged at 5000 rpm for 5 min. A 2-mL upper organic layer was taken and translocated to a d-SPE tube containing 600 mg MgSO_4_, 300 mg PSA and 100 mg alumina. The extract was shaken for 1 min and centrifuged. After centrifugation, 200 μL of upper layer was taken and placed into a vial for the analysis of pyrethroid residues using gas chromatography equipped with a micro electron capture detector (GC-μECD).

##### Roasted coffee powder and coffee sludge

2.5 g fine roasted coffee powder and 2.5 g coffee sludge were added with 10 ml acetonitrile and shaken for 30 sec. Then 4 g of MgSO_4_, 1 g of NaCl, 1 g of C_6_H_9_Na_3_O_9_ and 0.5 g C_6_H_5_Na_3_O_7_ were added and shaken for 30 sec. Samples were later centrifuged at 5000 rpm for 5 min. Two mL upper organic layer was taken and placed into d-SPE tube containing 600 mg magnesium sulfate anhydrous, 100 mg PSA and 100 mg alumina then shaken for 1 min and centrifuged thereafter at 5000 rpm for 5 min. After centrifugation, 200 μL of upper layer of individual sample was taken and placed into a vial for GC-μECD analysis.

#### Heavy metals

The acid digestion method was used to extract 9 metals: i.e., Cd, Co, Cr, Cu, Fe, Mn, Ni, Pb and Zn in all samples. Each metal concentration was determined using a flame atomic absorption spectrophotometer (FAAS). The digestion procedures of each sample type were as follows:

##### Green coffee beans, roasted coffee powder and coffee sludge

0.5 g of individual sample were digested with 5 mL mixture of 70% HNO_3_: 70% HClO_4_ (3:1 v/v) at 180°C for at least 3 h. The digested solution was maintained under this condition until clear solution indicating complete digestion was observed. Each sample was filtered using No. 4 Whatman filter paper (Germany) and then adjusted to 25 mL using ultra-high purity water and kept at 4°C until FAAS analysis.

##### Brew coffee drink

25 mL of each sample was digested with 5 mL 70% HNO_3_ at 180°C for at least 3 h. The digested samples were cooled to room temperature (25°C) and then filtered through No. 4 Whatman filter paper. The solution volume was adjusted to 25 mL using ultra-high purity water and kept at 4°C until FAAS analysis.

### Sample analysis

#### Gas chromatographic analysis

Pyrethroid residues analyses were carried out using an Agilent gas chromatography (GC 7890B, Agilent Technologies, Germany) with a micro-electron capture detector (μ+ ECD). The system was equipped with a 30 m x 0.32 mm x 0.25 μm fused silica capillary column (HP-5, Agilent Technologies, Germany). High purity (99.999%) helium and nitrogen were used as carrier gas and make up gas at a constant flow rate of 20 mL/min and 60 mL/min, respectively. The GC oven was operated as follows: initial temperature at 150°C held for 1 min, followed by the ramp of 15°C/min to 255°C and then 20°C/min to 300°C and held for 5 min. The injector and detector temperatures were set at 260°C and 315°C, respectively. The total run time was 28 min per sample. One microliter of each sample was injected to the GC system under splitless mode. Pyrethroid concentrations were quantified using standard calibration curves. Each sample was analyzed in triplicates to ensure reliable results.

The linearity, percentage recovery, precision, limit of detection (LOD) and limit of quantification (LOQ) for pyrethroid insecticides were determined under the European Commission (SANCO/12571/2013) criteria. The linearity (*r^2^* > 0.998) was validated for concentration range of 0.001 - 1.000 μg/g. The percentage recovery was determined using spiked known pyrethroid insecticide concentrations into each sample matrix. It was calculated by comparing the ratio of each spiked sample concentration to the analyzed concentration. The precision was obtained by spiking blank sample with each pyrethroid at 6 concentrations across the work range and express the variation in terms of relative standard deviation (%RSD). The LOD and LOQ were determined by comparing the target signal to noise ratio in blank sample spiked with known pyrethroid concentrations. For the LOD and LOQ the minimum signal to noise ratio was 3 and 10, respectively. The analytical performances were provided in Table 2.

**Table 2.** Results of percentage recovery (% recovery), percentage relative standard diviation (% RSD), limits of detection (LODs) and quatification (LOQs) of pyrethroid insecticides and metals.

| **Pyrethroid**<br>**insecticides/metals** | **% recovery** | **% RSD** | **LOD (µg/g)** | **LOQ (µg/g)** |
| --- | --- | --- | --- | --- |
| Cyfluthrin 1 | 86.37 | 9.27 | 0.005 | 0.030 |
| Cyfluthrin 2 | 98.62 | 11.25 | 0.004 | 0.017 |
| Cyfluthrin 3 | 79.29 | 12.02 | 0.003 | 0.010 |
| Cyfluthrin 4 | 102.65 | 12.30 | 0.005 | 0.017 |
| Flumethrin | 98.35 | 8.41 | 0.020 | 0.056 |
| Cypermethrin CisA | 97.95 | 11.47 | 0.003 | 0.021 |
| Cypermethrin CisB | 94.11 | 9.47 | 0.001 | 0.065 |
| Cypermethrin TransC | 88.87 | 11.78 | 0.005 | 0.019 |
| Cypermethrin TransD | 87.08 | 17.11 | 0.006 | 0.020 |
| Cd | 99.71 | 2.94 | 0.0005 | 0.002 |
| Co | 98.55 | 4.08 | 0.0003 | 0.001 |
| Cu | 99.81 | 3.43 | 0.001 | 0.003 |
| Cr | 98.04 | 3.91 | 0.0008 | 0.002 |
| Fe | 98.52 | 3.54 | 0.002 | 0.005 |
| Mn | 97.76 | 4.46 | 0.002 | 0.002 |
| Ni | 98.77 | 5.22 | 0.0004 | 0.001 |
| Pb | 99.51 | 2.16 | 0.002 | 0.005 |
| Zn | 97.76 | 7.12 | 0.0008 | 0.002 |

#### Heavy metal analysis

Nine heavy metal concentrations were determined using an air-acetylene FAAS (SpectrA 240B Agilent technologies, USA) equipped with deuterium background corrector. All metals were analyzed in an absorbance mode at the optimal wavelength for each metal as follows; Cd 228.8 nm, Co 240.7 nm, Cr 357.9 nm, Cu 324.8 nm, Fe 248.3 nm, Mn 279.5 nm, Ni 232.0 nm, Pb 217.0 nm, and Zn 213.9 nm. The individual metal concentration was calculated using its corresponding calibration curve.

The linearity (*r^2^* > 0.999) of calibration curve was evaluated using 5 replicates of 6 metal concentrations ranging from 0.05 - 2.00 μg/g for Cd, 0.05 - 5.00 μg/g for Co and Cu, 0.10 - 10.00 μg/g for Cr, 0.05 - 4.50 μg/g for Fe, 0.25 - 4.00 μg/g for Mn, 0.50 - 10.00 μg/g for Ni, 0.50 - 20.00 μg/g for Pb, and 0.10 - 2.00 μg/g for Zn. The accuracy of metal analysis was assessed using known spiked blank samples and the precision was also assessed by analyzing %RSD. The LOD and LOQ of each metal were calculated using signal-to-noise ratio (S/N) of 3 and 10, respectively. The obtained values were presented in Table 2.

### Determination of processing factor

The processing factor (PF) for each transformation step (roasting, brewing and sludge) was calculated as the ratio of pyrethroid insecticide or metal in processed samples (roasted coffee powder, brew coffee drink, coffee sludge) (μg/kg) to those in non-processed samples (green coffee bean) (μg/kg) using the following equation:

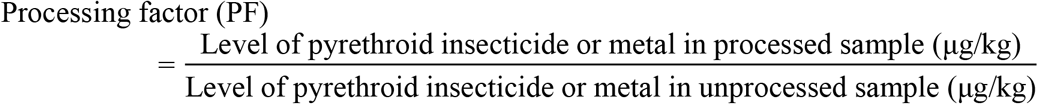

The PF of < 1 (reduction factor) indicated that there was a sample reduction of pyrethroid insecticide or metal by the processing step, whereas PF > 1 (concentration factor) indicated that there was no reduction in those toxic residues (16). In addition, the percentage reduction (% reduction) for individual processing step was calculated using the following equation:

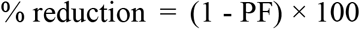

### Human health risk assessment

The estimate daily intake/EDI, health risk index/HRI (the ratio of calculated EDI to ADI or RfD) and target hazard quotients (THQs) were used to evaluate human health risk from insecticide-and metal-contaminated coffee drink. Risk determination was also calculated for the average consumption of brew coffee drink and at the 97.5^th^ percentiles of coffee drinker (extreme consumer). These allowed more comprehensive evaluation of human health risk associated with the consumption of pyrethroid and metal residues in coffee drink. The calculation formulae of EDI, HRI and THQ were listed as follows:

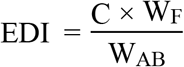

where C = pyrethroid or metal concentration in brew coffee sample (μg/mL); W_F_ = daily average consumption of eater only in Thailand (mL/person/day) (216.06 mL/person/day for normal and 330 mL/person/day for the 97.5^th^ percentile) and W_AB_ = average body weight (70 kg for adults).

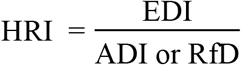

where ADI = the acceptable daily intake, RfD = oral reference dose (Σ cypermethrin 0.010 mg/kg bw/day, Σ cyfluthrin 0.004 mg/kg bw/day, Cd 0.001 mg/kg bw/day, Cr 0.003 mg/kg bw/day, Cu 0.500 mg/kg bw/day, Fe 0.800 mg/kg bw/day, Mn 0.140 mg/kg bw/day, Ni 0.020 mg/kg bw/day, Pb 0.025 mg/kg bw/day, and Zn 0.300 mg/kg bw/day)

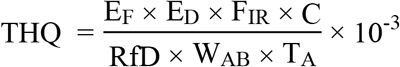

where E_F_ = exposure frequency (365 day/year), E_D_ = exposure duration (70 years), F_IR_ = food ingestion rate (mL/person/day) (216.06 mL/person/day for normal and 330 mL/person/day for the 97.5^th^ percentile), RfD = oral reference dose (mg/kg bw/day) (Σcyfluthrin 0.004 mg/kg bw/day, Cd 0.001 mg/kg bw/day, Cr 0.003 mg/kg bw/day, Cu 0.500 mg/kg bw/day, Fe 0.800 mg/kg bw/day, Mn 0.140 mg/kg bw/day, Ni 0.020 mg/kg bw/day, Pb 0.025 mg/kg bw/day, and Zn 0.300 mg/kg bw/day) (17, 18), W_AB_ = average body weight (70 kg for adults) and T_A_ = average exposure time (365 days/year x lifetime, assuming 70 years).

### Data analysis

Statistical analysis was performed using R statistical software version 3.4.3 (2017-11-30). The one-way Analysis of Variance (ANOVA) was used to test significant differences (*p* < 0.05) and similarities among insecticide residues found in coffee samples undergone different processes.

## Results and Discussion

### Pyrethroid residues in coffee samples

Residues of pyrethroid insecticides in green coffee beans, roasted coffee powder, brew coffee drink and coffee sludge are shown in Table 3. All studied pyrethorids were predominantly found in green coffee beans from various processes (dried, semi-washed, and washed) but not in brew coffee drink. The residues in green coffee beans differed significantly (*p* < 0.05) among processing steps (Table 3). Cyfluthrin 2, 3, cypermethrin CisA, and Σcyfluthrin concentrations in dried processed coffee beans were the highest (*p* < 0.05). In addition, concentrations of flumethrin and cypermethrin TransD found in semi-washed processed coffee beans were significantly the highest (*p* < 0.05). Simple washing process was proven to easily remove these residues (19). Pesticides and insecticides were removed from the outer and silver skins of the beans following washed and semi-washed processes (3). However, this residue removal may not always correlate to the water solubility of each residue compound. The peeling and refrigeration storage may also affect residue reduction (20).

**Table 3.** Average ± SE (μg/g or μg/mL) and percentage detection (%) of pyrethroid insecticide and metal concentrations in coffee samples from dried (D), semi-washed (SW) washed (W) processes.

| Pyrethroid<br>insecticides/metals | Average ± SE of pyrethroid insecticide and metal concentrations ($\mu\text{g/g}$ or $\mu\text{g/mL}$) and percentage detection (in parenthesis) | | | | | | | | | | | |
| --- | --- | --- | --- | --- | --- | --- | --- | --- | --- | --- | --- | --- |
|  | Green coffee beans |  |  | Roasted coffee powder |  |  | Brew coffee drink |  |  | Coffee sludge |  |  |
|  | D | SW | W | D | SW | W | D | SW | W | D | SW | W |
| Cyfluthrin 1 | 0.25±0.16<sup>a</sup><br>(66.67%) | 0.11±0.04<sup>a</sup><br>(60.00%) | 0.04±0.01<sup>a</sup><br>(50.00%) | 0.28±0.07<sup>b</sup><br>(26.67%) | 0.16±0.02<sup>ab</sup><br>(40.00%) | 0.02±0.00<sup>a</sup><br>(16.67%) | ND | ND | ND | 0.01±0.00<sup>a</sup><br>(26.67%) | 0.01±0.00<sup>a</sup><br>(40.00%) | 0.01±0.00<sup>a</sup><br>(20.00%) |
| Cyfluthrin 2 | 0.84±0.25<sup>b</sup><br>(100.00%) | 0.13±0.03<sup>a</sup><br>(60.00%) | 0.10±0.02<sup>a</sup><br>(90.00%) | ND | ND | ND | ND | ND | ND | ND | ND | ND |
| Cyfluthrin 3 | 0.19±0.04<sup>b</sup><br>(60.00%) | 0.04±0.02<sup>a</sup><br>(60.00%) | 0.03±0.01<sup>a</sup><br>(20.00%) | ND | ND | ND | ND | ND | ND | ND | ND | ND |
| Cyfluthrin 4 | 0.18±0.04<sup>a</sup><br>(86.67%) | 0.09±0.05<sup>a</sup><br>(60.00%) | 0.06±0.02<sup>a</sup><br>(33.33%) | ND | ND | ND | ND | ND | ND | ND | ND | ND |
| ΣCyfluthrin | 1.28±0.33<sup>b</sup><br>(100.00%) | 0.23±0.11<sup>a</sup><br>(100.00%) | 0.14±0.03<sup>a</sup><br>(93.33%) | 0.28±0.07<sup>b</sup><br>(26.67%) | 0.16±0.02<sup>ab</sup><br>(40.00%) | 0.02±0.00<sup>a</sup><br>(16.67%) | ND | ND | ND | 0.01±0.00<sup>a</sup><br>(26.67%) | 0.01±0.00<sup>a</sup><br>(40.00%) | 0.01±0.00<sup>a</sup><br>(20.00%) |
| Flumethrin | 0.39±0.06<sup>a</sup><br>(100.00%) | 0.75±0.01<sup>b</sup><br>(40.00%) | 0.24±0.04<sup>a</sup><br>(80.00%) | 0.06±0.00<sup>a</sup><br>(6.67%) | 0.06±0.02<sup>a</sup><br>(60.00%) | 0.06±0.03<sup>a</sup><br>(53.33%) | ND | ND | ND | 0.03±0.01<sup>a</sup><br>(26.67%) | ND | 0.02±0.01<sup>a</sup><br>(16.67%) |
| Cypermethrin CisA | 0.06±0.01<sup>b</sup><br>(60.00%) | 0.01±0.00<sup>a</sup><br>(20.00%) | 0.02±0.00<sup>a</sup><br>(6.67%) | ND | ND | ND | ND | ND | ND | ND | ND | ND |
| Cypermethrin CisB | 0.02±0.01<sup>a</sup><br>(33.33%) | 0.02±0.00<sup>a</sup><br>(20.00%) | 0.03±0.01<sup>a</sup><br>(66.67%) | 0.00<sup>a</sup><br>(6.67%) | 0.00<sup>a</sup><br>(20.00%) | 0.01±0.00<sup>a</sup><br>(43.33%) | ND | ND | ND | ND | ND | 0.001±0.00<sup>a</sup><br>(100.00%) |
| Cypermethrin<br>TransC | 0.01±0.00<sup>a</sup><br>(20.00%) | 0.01<sup>a</sup><br>(20.00%) | 0.01±0.00<sup>a</sup><br>(53.33%) | ND | ND | ND | ND | ND | ND | ND | ND | ND |
| Cypermethrin<br>TransD | 0.09±0.01<sup>ab</sup><br>(93.33%) | 0.11±0.05<sup>b</sup><br>(40.00%) | 0.06±0.01<sup>a</sup><br>(83.33%) | ND | 0.01±0.00<sup>a</sup><br>(60.00%) | 0.01±0.00<sup>a</sup><br>(56.67%) | ND | ND | ND | ND | 0.02±0.00<sup>a</sup><br>(20.00%) | 0.01±0.00<sup>a</sup><br>(20.00%) |
| ΣCypermethrin | 0.12±0.02<sup>a</sup><br>(100.00%) | 0.09±0.04<sup>a</sup><br>(60.00%) | 0.08±0.01<sup>a</sup><br>(93.33%) | 0.00<sup>a</sup><br>(6.67%) | 0.01±0.00<br>(60.00%) | 0.01±0.00<sup>a</sup><br>(33.33%) | ND | ND | ND | ND | 0.01±0.00<sup>a</sup><br>(20.00%) | 0.01±0.00<sup>a</sup><br>(26.67%) |
| Cd | 0.67±0.04<sup>ab</sup> | 0.64±0.02<sup>a</sup> | 0.74±0.02<sup>b</sup> | 0.50±0.03<sup>a</sup> | 0.56±0.01<sup>a</sup> | 0.56±0.02<sup>a</sup> | 0.01±0.00<sup>b</sup> | ND | 0.01±0.00<sup>a</sup> | 0.87±0.30<sup>a</sup> | 0.44±0.04<sup>a</sup> | 0.48±0.02<sup>a</sup> |

| Pyrethroid<br>insecticides/metals | Average ± SE of pyrethroid insecticide and metal concentrations (µg/g or µg/mL) and percentage detection (in parenthesis) |  |  |  |  |  |  |  |  |  |  |  |
| --- | --- | --- | --- | --- | --- | --- | --- | --- | --- | --- | --- | --- |
|  | Green coffee beans |  |  | Roasted coffee powder |  |  | Brew coffee drink |  |  | Coffee sludge |  |  |
|  | D | SW | W | D | SW | W | D | SW | W | D | SW | W |
|  | (100.00%) | (100.00%) | (100.00%) | (100.00%) | (100.00%) | (100.00%) | (100.00%) |  | (50.00%) | (100.00%) | (100.00%) | (100.00%) |
| Co | 0.88±0.10<sup>a</sup> | 1.32±0.43<sup>a</sup> | 0.92±0.11<sup>a</sup> | 1.87±0.14<sup>a</sup> | 1.91±0.28<sup>a</sup> | 1.89±0.14<sup>a</sup> | 0.02±0.01<sup>a</sup> | 0.01±0.01<sup>a</sup> | 0.02±0.00<sup>a</sup> | 1.35±0.17<sup>a</sup> | 1.19±0.16<sup>a</sup> | 1.45±0.11<sup>a</sup> |
|  | (73.33%) | (80.00%) | (96.67%) | (100.00%) | (100.00%) | (100.00%) | (80.00%) | (40.00%) | (80.00%) | (100.00%) | (100.00%) | (100.00%) |
| Cu | 35.79±16.66<sup>a</sup> | 42.56±18.12<sup>a</sup> | 22.00±5.23<sup>a</sup> | 12.65±2.89<sup>a</sup> | 12.41±1.69<sup>a</sup> | 15.19±1.60<sup>a</sup> | 0.02±0.00<sup>b</sup> | 0.02±0.01<sup>ab</sup> | 0.01±0.00<sup>a</sup> | 12.03±1.73<sup>a</sup> | 15.21±3.00<sup>a</sup> | 14.33±1.54<sup>a</sup> |
|  | (100.00%) | (100.00%) | (100.00%) | (100.00%) | (100.00%) | (100.00%) | (100.00%) | (100.00%) | (100.00%) | (100.00%) | (100.00%) | (100.00%) |
| Cr | ND | ND | ND | 0.13±0.00<sup>b</sup> | 0.97±0.19<sup>a</sup> | 0.55±0.12<sup>ab</sup> | 0.03±0.00<sup>a</sup> | 0.04±0.00<sup>a</sup> | 0.03±0.00<sup>a</sup> | 0.20±0.04<sup>a</sup> | 0.26±0.06<sup>a</sup> | 0.37±0.08<sup>a</sup> |
|  |  |  |  | (13.33%) | (80.00%) | (40.00%) | (100.00%) | (100.00%) | (100.00%) | (20.00%) | (100.00%) | (46.67%) |
| Fe | 34.74±3.78<sup>a</sup> | 27.76±2.29<sup>a</sup> | 27.11±0.98<sup>a</sup> | 32.67±2.33<sup>b</sup> | 32.07±2.75<sup>ab</sup> | 27.07±0.45<sup>a</sup> | 0.15±0.01<sup>b</sup> | 0.14±0.02<sup>b</sup> | 0.11±0.01<sup>a</sup> | 39.72±5.23<sup>a</sup> | 34.97±2.66<sup>a</sup> | 28.78±0.76<sup>a</sup> |
|  | (100.00%) | (100.00%) | (100.00%) | (100.00%) | (100.00%) | (100.00%) | (100.00%) | (100.00%) | (100.00%) | (100.00%) | (100.00%) | (100.00%) |
| Mn | 20.40±1.99<sup>ab</sup> | 19.61±3.40<sup>a</sup> | 30.56±2.39<sup>b</sup> | 17.70±1.65<sup>a</sup> | 21.35±4.13<sup>ab</sup> | 31.21±2.36<sup>b</sup> | 0.34±0.06<sup>a</sup> | 0.33±0.04<sup>a</sup> | 0.62±0.09<sup>a</sup> | 19.09±1.31<sup>a</sup> | 20.11±4.57<sup>a</sup> | 27.45±2.07<sup>a</sup> |
|  | (100.00%) | (100.00%) | (100.00%) | (100.00%) | (100.00%) | (100.00%) | (100.00%) | (100.00%) | (100.00%) | (100.00%) | (100.00%) | (100.00%) |
| Ni | 7.67±0.60<sup>b</sup> | 10.41±0.60<sup>a</sup> | 9.70±0.54<sup>ab</sup> | 4.70±0.38<sup>a</sup> | 4.28±0.53<sup>a</sup> | 3.71±0.23<sup>a</sup> | 0.07±0.01<sup>b</sup> | 0.05±0.02<sup>ab</sup> | 0.04±0.01<sup>a</sup> | 4.26±0.47<sup>a</sup> | 4.02±0.97<sup>a</sup> | 3.73±0.27<sup>a</sup> |
|  | (100.00%) | (100.00%) | (100.00%) | (100.00%) | (100.00%) | (96.67%) | (100.00%) | (80.00%) | (83.33%) | (100.00%) | (100.00%) | (100.00%) |
| Pb | 20.14±2.16<sup>a</sup> | 20.71±2.52<sup>a</sup> | 16.92±0.75<sup>a</sup> | 23.32±0.90<sup>a</sup> | 23.44±0.98<sup>a</sup> | 21.19±0.54<sup>a</sup> | 0.28±0.02<sup>ab</sup> | 0.26±0.02<sup>a</sup> | 0.31±0.01<sup>b</sup> | 18.02±0.67<sup>a</sup> | 18.02±0.43<sup>a</sup> | 17.16±0.40<sup>a</sup> |
|  | (100.00%) | (100.00%) | (100.00%) | (100.00%) | (100.00%) | (100.00%) | (100.00%) | (100.00%) | (100.00%) | (100.00%) | (100.00%) | (100.00%) |
| Zn | 20.84±7.19<sup>a</sup> | 30.13±11.61<sup>a</sup> | 15.59±3.46<sup>a</sup> | 9.27±1.65<sup>a</sup> | 7.66±0.92<sup>a</sup> | 8.64±0.96<sup>a</sup> | 0.07±0.01<sup>a</sup> | 0.07±0.01<sup>a</sup> | 0.08±0.01<sup>a</sup> | 8.50±0.97<sup>a</sup> | 9.84±1.75<sup>a</sup> | 7.49±0.93<sup>a</sup> |
|  | (100.00%) | (100.00%) | (100.00%) | (100.00%) | (100.00%) | (100.00%) | (100.00%) | (100.00%) | (100.00%) | (100.00%) | (100.00%) | (100.00%) |

Means followed by different letters indicated significant ( $p < 0.05$ ) differences among coffee bean processes.
Nd : Non detectable (concentrations < LOD)

Concentrations of all studied pyrethroids in brew coffee drink (Table 3) were lower than their corresponding limits of detection (Table 2). The roasting and brew coffee steps reduced pyrethroid levels in coffee drink. Decreases in the amount of pyrethroids were via molecular structure breakdown (21) and physico-chemical properties e.g., evaporation, co-distillation and thermal degradation which may vary with chemical nature of the individual compound (22). There were no statistical differences in pyrethroid concentrations (*p* > 0.05) among coffee bean processes in roasted coffee beans and brew coffee drinks.

Most pyrethroid residue levels and their detection percentages in coffee sludge samples were lower than those found in roasted coffee beans (Table 3). Cyfluthrin 1 concentration found in coffee sludge sample from semi-washed beans was the highest (0.16 ± 0.02 μg/g). However, all pyrethroid residue concentrations in coffee sludge did not significantly differ among coffee bean processes (*p* > 0.05).

### Heavy metal residues in coffee samples

Most of the studied 9 metals were found in green coffee beans, roasted coffee powder, brew coffee drink, and coffee sludge except for Cr in green coffee beans and Cd in brew coffee drink (only in semi-washed process samples) (Table 3). In addition, the lowest and the highest metal concentrations were in brew coffee drink and green coffee beans, respectively. The significant differences (*p* > 0.05) of Cd and Fe were found among the brew coffee drink.

The highest trace and toxic metal concentrations in green coffee beans were Cu (42.56 ± 18.12 μg/g) and Pb (20.71 ± 2.52 μg/g) from semi-washed process. Copper is an inactive ingredient of some pesticides used in coffee culture. Its highest concentration was found in coffee bean sample from Ethiopia with the concentration approximately 4 times higher than those in previous reports (14, 15). Besides Cu, green coffee beans were contaminated with Pb at higher concentration than those reported in other studies (14, 15, 23). The metal concentration differences depend on cultivation areas characterized by different organoleptic features and chemical compositions, which could be used as a tool for characterization coffee variation (12, 24). The Pb residue can affect the neurological system resulting in central and peripheral nervous system damages. It can also interfere with vitamin D conversion and calcium homeostasis, inhibit hemoglobin synthesis, as well as be a potential carcinogen (25). While excess Cu level may alter the metabolic functions and cause Menkes and Wilson’s disease (26). Organic fertilizers, organic manures and pesticides used in cultivation usually were sources of different metal contaminations (12). However, there was no regulation of each metal residue in green coffee beans.

No significant differences (*p* > 0.05) of metal residue concentrations in roasted coffee powder existed among coffee bean processes (Table 3). This indicated that the roasting process affected the metal concentrations in coffee beans. Mn concentration (32.21 ± 2.36 μg/g) was the highest in washed process whereas Fe concentrations (32.67 ± 2.33 μg/g, 32.07 ± 2.75 μg/g) were the highest in dried and semi-washed processes in roasted powder samples, respectively. Similar Mn and Fe concentrations in roasted coffee beans were previously reported (27). Metal concentrations in roasted coffee powder were higher than those in green coffee beans. It is likely that high temperature (up to 250°C) used in the roasting process affected chemical compositions as well as biological activities of the green coffee beans (27). Increased metal concentrations, thus, were apparent (15, 27, 28). Not only the coffee bean origins and qualities but also the coffee bean processes could alter concentrations of both trace and toxic metals. The Pb concentrations in roasted coffee beans found in this study were above the EU permitted limit (0.2 mg/kg) (29).

The statistical differences of heavy metal concentrations (*p* > 0.05) among bean processes were found for Cd, Cu, Fe, Ni, and Pb in brew coffee drink. Most metals were detected except for Cd in semi-washed process. Concentrations of Mn (0.62 ± 0.09 μg/mL in washed process) and Pb (0.28 ± 0.02 μg/mL in dried process) were the highest among trace and toxic metals, respectively. The detected Mn concentrations were similar to the report of Nędzarek, Tórz (30) but slightly higher than those found in others. These differences were caused from a wide range of geographical areas in coffee culture (12, 24, 27).

There were no significant differences (*p* > 0.05) of metal concentrations in coffee sludge among bean processes. The metal found at the highest concentration was Fe (39.72 ± 5.23 μg/g) in dried process coffee sludge. Coffee sludge has currently been used by several means, e.g., organic fertilizer for agricultural crops including coffee, raw materials for bioethanol and biogas production (31). If metal concentrations are high like those found in this study, the transfer of toxic contaminants can be expected. Metal contaminated coffee sludge for such purposes should be of concern. The regulation of metal residues in coffee sludge should then be established.

### Determination of processing factors

The processing factor (PF) and percentage reduction (%) for individual pesticide and metal from each process were determined (Table 4). No significant difference (*p* > 0.05) of PF was found among the bean processes. All PFs of pyrethroids from all coffee bean processes were less than 1. This indicated that coffee roasting and brewing decreased pyrethroid residues. The results are in agreement with other studies whose PFs were decreased following food processing (3, 16). The roasting process plays the most important role in pyrethoriod reduction. However, the PF was not calculated for the pyrethroids not detected in samples. On the contrary, higher concentrations of most metals were observed after roasting indicated by PF greater than 1 (Table 4). The increased concentrations were likely from the evaporation of water and/or combustible mass in the coffee grain which concentrate metals in the coffee beans (27). The brew coffee drinks showed the opposite processing factors (PF < 1) with percentage reductions greater than 96%. This indicated that coffee brewing resulted in the reduction of all metal concentrations and were likely due to dilution effect as their weight reduction.

**Table 4.** Processing factor (PF) and percentage reduction (%) in coffee samples from dried, semi-washed and washed processes.

| Pyrethroid /<br>metals residues | Processing factor (PF) and percentage reduction (%) |  |  |  |  |  |  |  |  |
| --- | --- | --- | --- | --- | --- | --- | --- | --- | --- |
|  | Dried process |  |  | Semi-washed process |  |  | Washed process |  |  |
|  | Roasted<br>beans | Brew coffee<br>drink | Coffee<br>sludge | Roasted<br>beans | Brew coffee<br>drink | Coffee<br>sludge | Roasted<br>beans | Brew coffee<br>drink | Coffee<br>sludge |
| **Cyfluthrin 1** | 0.56<br>(43.77%) | NA | 0.01<br>(99.06%) | 0.00<br>(100.00%) | NA | 0.00<br>(100.00%) | 0.01<br>(99.19%) | NA | 0.12<br>(88.08%) |
| **Cyfluthrin 2** | 0.00<br>(100.00%) | NA | 0.00<br>(100.00%) | 0.00<br>(100.00%) | NA | 0.00<br>(100.00%) | 0.00<br>(100.00%) | NA | 0.00<br>(100.00%) |
| **Cyfluthrin 3** | 0.00<br>(100.00%) | NA | 0.00<br>(100.00%) | 0.00<br>(100.00%) | NA | 0.00<br>(100.00%) | 0.00<br>(100.00%) | NA | 0.00<br>(100.00%) |
| **Cyfluthrin 4** | 0.00<br>(100.00%) | NA | 0.00<br>(100.00%) | 0.00<br>(100.00%) | NA | 0.00<br>(100.00%) | 0.00<br>(100.00%) | NA | 0.00<br>(100.00%) |
| **ΣCyfluthrin** | 0.59<br>(40.83%) | NA | 0.01<br>(99.81%) | 0.28<br>(72.21%) | NA | 0.01<br>(98.81%) | 0.02<br>(98.44%) | NA | 0.06<br>(94.50%) |
| **Flumethrin** | 0.02<br>(98.05%) | NA | 0.04<br>(96.00%) | 0.07<br>(93.16%) | NA | 0.00<br>(100.00%) | 0.27<br>(73.26%) | NA | 0.01<br>(99.14%) |
| **Cypermethrin**<br>**CisA** | 0.00<br>(100.00%) | NA | 0.00<br>(100.00%) | 0.00<br>(100.00%) | NA | 0.00<br>(100.00%) | 0.00<br>(100.00%) | NA | 0.00<br>(100.00%) |
| **Cypermethrin**<br>**CisB** | 0.04<br>(95.64%) | NA | 0.00<br>(100.00%) | 0.05<br>(94.66%) | NA | 0.00<br>(100.00%) | 0.10<br>(89.62%) | NA | 0.01<br>(98.68%) |
| **Cypermethrin**<br>**TransC** | 0.00<br>(100.00%) | NA | 0.00<br>(100.00%) | 0.00<br>(100.00%) | NA | 0.00<br>(100.00%) | 0.00<br>(100.00%) | NA | 0.00<br>(100.00%) |
| **Cypermethrin**<br>**TransD** | 0.00<br>(100.00%) | NA | 0.00<br>(100.00%) | 0.12<br>(88.39%) | NA | 0.00<br>(100.00%) | 0.05<br>(95.14%) | NA | 0.01<br>(98.76%) |
| **ΣCypermethrin** | 0.01<br>(99.85%) | NA | 0.00<br>(100.00%) | 0.11<br>(89.27) | NA | 0.00<br>(100.00%) | 0.97<br>(90.26%) | NA | 0.05<br>(95.32%) |
| **Cd** | 0.79<br>(21.54%) | 0.01<br>(98.99%) | 1.37<br>(-36.89%) | 0.88<br>(12.09%) | NA | 0.69<br>(30.89%) | 0.77<br>(22.77%) | 0.00<br>(99.78%) | 0.65<br>(35.26%) |
| **Co** | 2.36<br>(-135.67) | 0.02<br>(97.80%) | 1.65<br>(-65.08%) | 2.21<br>(-121.42%) | 0.01<br>(99.74%) | 1.39<br>(-39.25%) | 4.57<br>(-356.99%) | 0.04<br>(96.230%) | 3.32<br>(-232.00%) |
|  | Dried process |  |  | Semi-washed process |  |  | Washed process |  |  |
|  | Roasted<br>beans | Brew coffee<br>drink | Coffee<br>sludge | Roasted<br>beans | Brew coffee<br>drink | Coffee<br>sludge | Roasted<br>beans | Brew coffee<br>drink | Coffee<br>sludge |
| **Cr** | NA | NA | NA | NA | NA | NA | NA | NA | NA |
| **Cu** | 1.04<br>(-4.408%) | 0.00<br>(99.79%) | 1.08<br>(-7.76%) | 0.68<br>(32.11%) | 0.01<br>(99.92%) | 1.21<br>(-20.92%) | 1.19<br>(-18.59%) | 0.01<br>(99.88%) | 1.11<br>(-10.53%) |
| **Fe** | 1.03<br>(-3.27%) | 0.01<br>(99.54%) | 1.25<br>(-25.34%) | 1.16<br>(-16.20%) | 0.01<br>(99.50%) | 1.27<br>(-26.88%) | 1.02<br>(-2.16%) | 0.01<br>(99.58%) | 1.10<br>(-9.06%) |
| **Mn** | 0.88<br>(11.69%) | 0.02<br>(98.45%) | 0.98<br>(2.16%) | 1.11<br>(-10.85%) | 0.02<br>(98.23%) | 1.03<br>(-2.97%) | 1.05<br>(-4.68) | 0.02<br>(98.06%) | 0.93<br>(7.07%) |
| **Ni** | 0.97<br>(2.64%) | 0.02<br>(98.24%) | 0.86<br>(14.55%) | 0.41<br>(59.29%) | 0.01<br>(99.51%) | 0.38<br>(61.56%) | 0.39<br>(60.90%) | 0.014<br>(99.60%) | 0.41<br>(59.55%) |
| **Pb** | 1.28<br>(-28.42%) | 0.02<br>(98.42%) | 1.00<br>(0.20%) | 1.20<br>(-20.35%) | 0.013<br>(98.67%) | 0.91<br>(8.80%) | 1.31<br>(-31.33%) | 0.029<br>(98.06%) | 1.06<br>(-5.75%) |
| **Zn** | 0.99<br>(1.43%) | 0.01<br>(99.35%) | 0.86<br>(13.88%) | 0.50<br>(49.56%) | 0.01<br>(99.48%) | 0.77<br>(23.35%) | 1.23<br>(-22.85%) | 0.01<br>(99.05%) | 1.10<br>(-9.70%) |

NA (not available): the processing factor was not calculated due to residue of pesticides or metals was not detected.

### Human health risk assessment

Human health risks from pyrethroid insecticide and metal contamination in brew coffee drink were assessed by considering two consumptions scenario, i.e., normal consumption and extreme consumption approaches (97.5^th^ percentile) of Thai population. The estimate daily intakes (EDIs) and their health risk indices (HRIs) were presented in Table 5. None of pyrethroid pesticides were detectable in brew coffee drinks whereas all metals were found but the concentrations were lower than reference ADIs or RfDs (HRI < 1). The EDIs have shown that toxic metals (Cd, Cr, Ni and Pb) and pyrethroid insecticides (cyfluthrin, cypermethrin and flumethrin) contaminations in brew coffee drinks are safe for both normal and the 97.5^th^ percentiles of Thai population. Concentration of Pb found in brew coffee drink (0.31 μg/mL) was the highest among toxic metals. Its highest EDIs via normal consumption (216.06 mL/day) and the 97.5^th^ percentile (330 mL/day) consumption for Thai coffee drinker (32) were 0.97 mg/kg bw/day (washed process) and 1.48 mg/kg bw/day (washed process), respectively. In addition, all calculated HRIs which were less than 1 (Table 5) indicated that human consumption of all studied samples was relatively safe. Trace metals such as Co, Cu, Fe, Mn, Ni and Zn are cofactors of a large number of enzymes. Their trace but adequate amounts are essential for normal body functions (Cu 0.9 mg/day; Fe 8-18 mg/day; Mn 1.8-2.3 mg/day; Ni 0.5 mg/day and Zn 8-11 mg/day) (33). The excess amount of these trace metals, however, could cause adverse human health effects. The estimated consumptions of these metals in brew coffee drink from this study were higher than normal human requirement but still safe for the consumption. A few studies showed no health risk effects of mineral intake following coffee consumption similar to this study (14, 27, 34). The THQ values were calculated and used to assess for human health risk for long-time consumers (Table 6). In this study, the THQs of all investigated analytes were less than 1. The consumption of brew coffee drinks from all studied coffee bean samples, thus, are unlikely to pose adverse effects on human health.

**Table 5.**
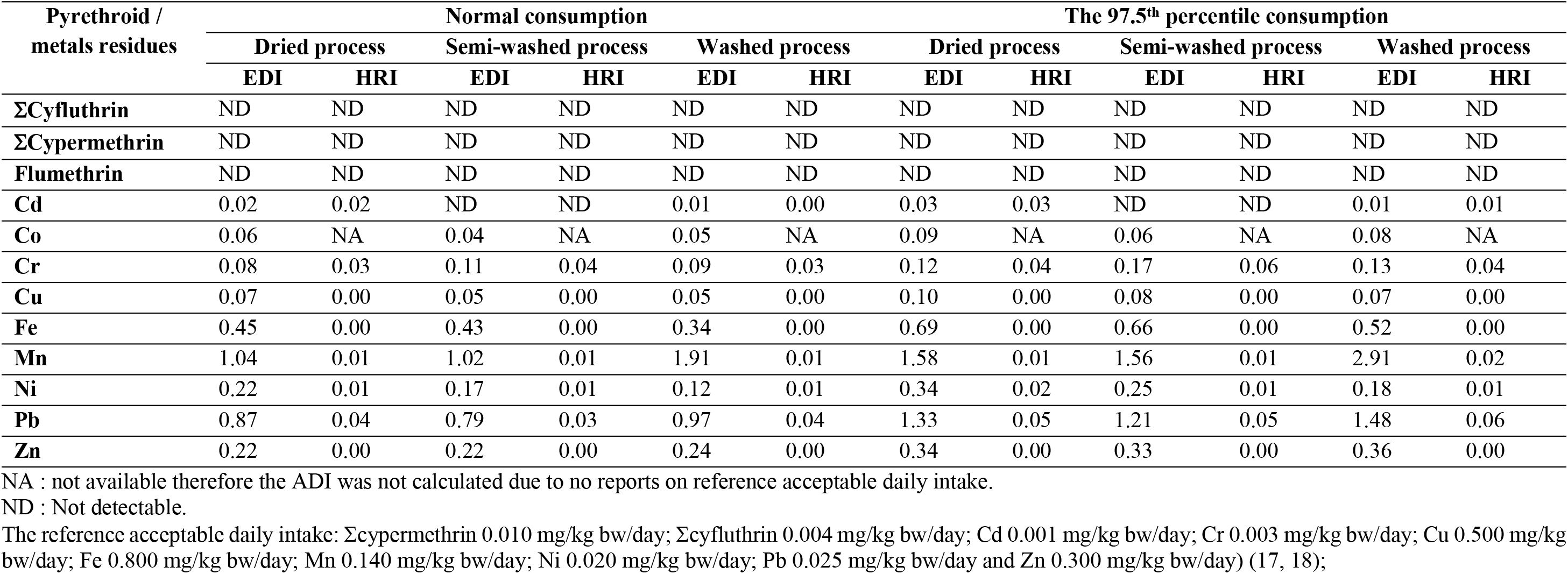
Estimated normal dietery (EDI) of exposure levels to residues of pesticides (μg/kg bw/day) and metals (μg/kg bw/day) detected in brew coffee drink and Health Risk Index (HRI) based on normal consumption and the 97^th^ percentile consumption of Thai population.

**Table 6.** Average target hazard quotient (THQ) with the consumption of brew coffee drink based on normal consumption and the 97^th^ percentile consumption of Thai population.

| Pyrethroid /<br>metals residues | Target hazard quotient (THQ) |  |  |  |  |  |
| --- | --- | --- | --- | --- | --- | --- |
|  | Normal consumption |  |  | The 97.5<sup>th</sup> percentile consumption |  |  |
|  | Dried process | Semi-washed process | Washed process | Dried process | Semi-washed process | Washed process |
| **ΣCyfluthrin** | ND | ND | ND | ND | ND | ND |
| **ΣCypermethrin** | ND | ND | ND | ND | ND | ND |
| **Flumethrin** | ND | ND | ND | ND | ND | ND |
| **Cd** | 0.02 | ND | 0.01 | 0.03 | ND | 0.01 |
| **Co** | NA | NA | NA | NA | NA | NA |
| **Cr** | 0.03 | 0.05 | 0.04 | 0.04 | 0.06 | 0.04 |
| **Cu** | <0.01 | <0.01 | <0.01 | <0.01 | <0.01 | <0.01 |
| **Fe** | <0.01 | <0.01 | <0.01 | 0.01 | 0.01 | 0.01 |
| **Mn** | 0.01 | 0.01 | 0.02 | 0.01 | 0.01 | 0.02 |
| **Ni** | 0.01 | 0.01 | 0.01 | 0.02 | 0.01 | 0.01 |
| **Pb** | 0.04 | 0.04 | 0.05 | 0.05 | 0.05 | 0.06 |
| **Zn** | <0.01 | <0.01 | <0.01 | 0.01 | 0.01 | 0.01 |

NA : the THQ was not calculated due to no report on reference acceptable daily intake. ND : Not detectabl

## Conclusion

Coffee is among the most popular beverages in all countries. Insecticides and metals introduced during coffee cultures could result in their contaminations in the coffee beans. These contaminants could be reduced during coffee bean processing including dried, semi-washed, and washed processes. In this study, 3 pyrethroid insecticides and 9 metals were determined in coffee samples, i.e., green coffee beans, roasted coffee powder, brew coffee drink, and coffee sludge. Coffee roasting and brewing could reduce the concentrations of both investigated pyrethroid insecticides and metals indicated by the PF values which were less than Based on the normal and the 97.5^th^ percentile consumption of Thai population, it can be concluded that the consumption of brew coffee drink is unlikely to pose adverse effects on human health. Uses of coffee by-product (coffee sludge) as fertilizers or biogas raw materials should be of concerns since it may contain toxic contaminants.

## Acknowledgements

This research is supported in part by the Graduate Program Scholarship from the Graduate School, Kasetsart University.

